# Cancer Evolvability Determines Therapy Outcomes

**DOI:** 10.64898/2026.05.06.723143

**Authors:** Ranjini Bhattacharya, Anuraag Bukkuri, Robert A. Gatenby, Joel S. Brown

## Abstract

Cancer progression following treatment failure is an evolutionary process in which therapy acts as a selection pressure driving Darwinian selection on heritable variation to favor resistant clones. This ability to generate variation, i.e., the cancer’s evolvability, is a key determinant of how rapidly tumors adapt to therapy. Here, we present an evolutionary game-theoretic model to evaluate how evolvability shapes resistance dynamics under two treatment modalities: targeted therapy and chemotherapy.

We first compare cancer populations with fixed evolvabilities: low or high. Targeted therapy imposes a steep selection gradient, enabling rapid resistance evolution, while chemotherapy exerts a flatter gradient but drives tumors toward more extreme resistance strategies. We show that targeted therapy works better in low-evolvability cancers, whereas chemotherapy better controls high-evolvability populations. We then extend the model to incorporate facultative evolvability in which cancer cells dynamically adjust their evolvability in response to therapy-induced stress in which cells fine-tune the trade-off between acquiring higher resistance and limiting the costs of resistance and evolvability. The latter strategy sustains a higher tumor burden than fixed-evolvability populations.

To address the challenges of facultative evolvability for therapy efficacy, we develop and simulate an evolutionary double bind using sequential cycles of chemotherapy and targeted therapy. With an appropriate sequence and timing, this strategy can drive cancer cells with facultative evolvability to extinction. Our results highlight the importance of evolvability in shaping treatment response and underscore the need to incorporate evolutionary principles into therapy design.

## Introduction

Cancer is a disease shaped by evolutionary forces acting within the body. Cancer cells are constantly exposed to selection pressures in the tumor microenvironment as they compete for nutrients and space while avoiding host response. Cancer treatment represents an additional evolutionary selection pressure [1,2]. Natural selection favors cells that survive and proliferate under these microenvironmental conditions, leading to the emergence of fitter variants [3]. In the face of therapy, most cancer cells sensitive to therapy die. However, some cancer cells acquire traits that make them resistant to the drug, making subsequent treatment ineffective [4]. This often leads to relapse in patients after months or years in remission [5]. Evolution is, therefore, the main cause of treatment failure and subsequent disease progression.

Darwinian evolution requires: heritable variation, a struggle for survival, and a link between the two [6]. In the context of cancer therapy, drug administration creates a strong selection pressure, i.e., a struggle for survival, within the tumor. Only those variants capable of surviving in this drug-exposed environment are selected by natural selection [7]. Resistance can be either intrinsic or acquired [5,8]. Intrinsic resistance arises when resistant subclones exist at low frequencies within the tumor prior to drug administration and expand after therapy eliminates sensitive cells. For instance, the presence of HER2-overexpressing cells before treatment is associated with poor response to cisplatin therapy [9]. Acquired resistance, on the other hand, emerges during treatment as cancer cells develop new resistance traits. A common mechanism involves the upregulation of drug efflux pumps through increased expression of ABC transporters following therapy exposure [10,11] conferring multidrug resistance to diverse therapies [12].

The rate at which a cancer cell population evolves is influenced by the strength of selection pressures and the amount of heritable variation it generates via mutations, epigenetic changes, copy number variation, chromosomal rearrangements, polyploidy, etc. [6,13]. The capacity to generate heritable variation determines the population’s evolvability [7,14]. We illustrate this using the adaptive landscape shown in Fig. 1 [15,16]. An adaptive landscape maps the fitness of a focal cell introduced into a population with a specific density distribution and composition of trait values. It illustrates both the current fitness of the population given its existing strategy and the optimal strategy under the prevailing environmental conditions. The adaptive landscape shown represents a snapshot in time and could change as the population evolves. The slope of the landscape represents the selection gradient, and the step size the population can take in either direction represents its evolvability [17].

**Figure 1:**
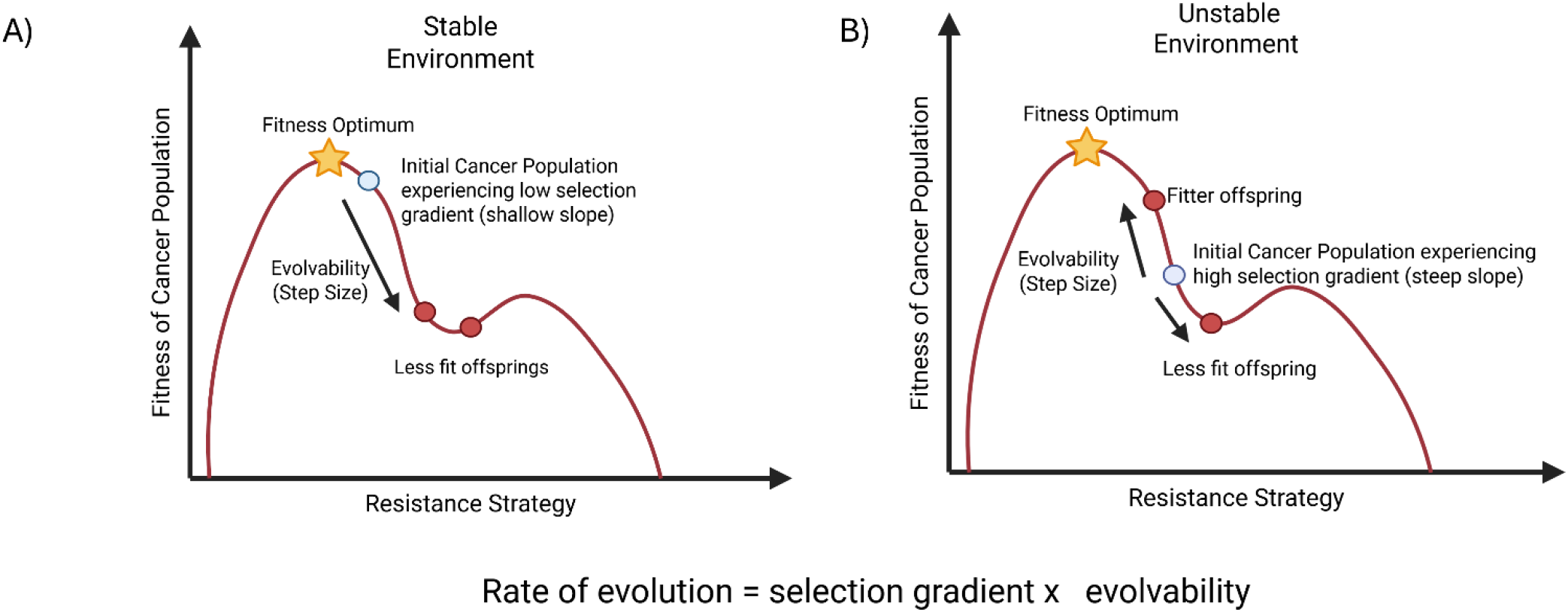
A Hypothetical Adaptive Landscape: The adaptive landscape maps trait value of a focal cell introduced into a population with a certain composition of traits. The slope of the landscape represents the selection gradient, and the step size a population can take along the landscape represents its evolvability (amount of heritable variation it can generate). The rate of evolution is proportional to the product of these two quantities. (A) Closer to the fitness optimum (Stable Environment): The selection gradient is shallow. A high evolvability in this environment could lead to the generation of less fit offsprings, which can reduce fitness and, in extreme cases, lead to mutational meltdown. (B) Closer to the fitness optimum (Unstable Environment): The population experiences a steep selection gradient. A high evolvability in such an environment is more likely to generate fitter offspring, accelerating adaptation toward the optimum. (Created using Biorender.com).

Biological pathways that regulate mutation rates [18,19], epigenetic reprogramming [20,21], mRNA translation [22,23], and phenotypic plasticity [24] all contribute to a population’s evolvability by altering the phenotypes of daughter cells. In the presence of therapy (catastrophically altered, unstable environment), high evolvability could speed up adaptation by generating resistant daughter cells (Fig. 1B) [25]. However, in a stable environment, where the population exists closer to the fitness optimum, high evolvability results in an excess of less fit individuals. However, in the extreme, an excessive rate of generating heritable variation could lead to a mutational meltdown [25–27] resulting in decrease fitness. Recently, Weh *et al*. modeled evolvability by explicitly including high and low mutation rates in an agent-based model. They showed how high mutation rates favor evolutionary rescue while preventing the population from evolving to its fitness optimum [28].

Cancer drug therapies include diverse mechanisms to induce cancer cell death, including cytotoxic treatments like chemotherapy [29] and targeted therapies [30,31]. Chemotherapy typically involves a combination of cytotoxic drugs that interfere with biological processes such as DNA replication or cell division. These agents broadly affect all cells regardless of their phenotype and are associated with significant off-target toxicity [32]. For instance, cisplatin forms DNA crosslinks that inhibit replication and transcription in both normal cells and cancer cells [33]. Because chemotherapy targets processes involved in cell cycling, it imposes a selective pressure that drives the emergence of higher and more heterogeneous resistance strategies. In contrast, targeted therapies are designed to inhibit specific molecular pathways that are implicated in cancer progression. These therapies are generally less toxic, with fewer off-target effects, but are effective only on cells that rely on the targeted oncogenic driver [34]. A well-known example is tamoxifen, which binds competitively to the estrogen receptor and blocks estrogen-mediated signaling in ER-positive breast cancer [35,36]. While initially effective, targeted therapies are often vulnerable to resistance, as cancer cells need only activate an alternative signalling pathway to bypass the inhibited one. For example, resistance to tamoxifen can develop through upregulation of growth factor signalling pathways, such as HER2 or IGF1R, which can activate downstream proliferative signals independent of estrogen receptor activity [37].

Here, we develop an evolutionary game-theoretic model to investigate how evolvability influences a cancer population’s response to chemotherapy and targeted therapy. A key distinction between these therapies is that chemotherapy is broadly effective across a wide range of cancer phenotypes but lacks specificity. In contrast, targeted therapy is highly specific, acting on a narrow subset of cancer cells. This makes targeted therapies more susceptible to resistance. We compare the therapeutic responses of three cancer population types: those with a constant low evolvability, constant high evolvability, and facultative evolvability. Facultative evolvability refers to a population’s ability to modulate the generation of heritable variation based on environmental cues, such as the presence or absence of stressors, such as treatment [38,39]. A growing body of research suggests that environmental conditions can influence evolvability; for instance, stress in the tumor microenvironment has been shown to increase mutagenesis, thereby enhancing evolvability [40,41].

We show that the three different populations respond differently to therapy. Based on the model’s predictions, we offer therapeutic recommendations tailored to the cancer population’s evolvability, with the goal of controlling the evolution of resistance and optimizing therapy outcomes.

## Methods

### Single Therapy Model

To investigate how evolvability influences the cancer population’s response to therapy, we develop a G-function model, a framework rooted in evolutionary game theory (EGT) [42,43]. EGT enables the study of how interactions between cancer cells and the tumor microenvironment shape their evolutionary trajectories [44]. The G-function framework, specifically, couples ecological dynamics (cell density) and evolutionary dynamics (trait distribution). At the heart of this approach is a fitness-generating function that defines how a population’s fitness depends on both its traits and ecological context. In our model, we focus on how the fitness of a cancer cell population is shaped by drug efficacy and the resistance trait [45]. We model resistance as a continuous trait instead of a discrete state. Equation 1 shows the G-function (*G*(*v, x*)) as a function of cell density (*x*) and the resistance strategy (*v*):

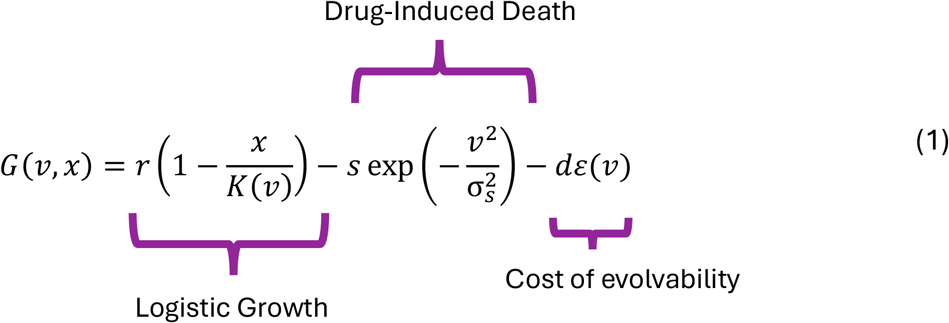

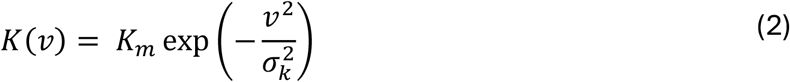

The first term in Equation 1 represents logistic growth, where *r* is the intrinsic per capita growth rate of the cancer cells and *K*(*v*) is the carrying capacity, modeled as a function of the resistance trait *v* ∈ [0, ∞) (Equation 2). The maximum carrying capacity occurs when *v* = 0 (no resistance) and decreases in a Gaussian manner as *v* deviates from 0. σ_*k*_ represents this deviation. This formulation imposes an ecological cost of resistance, such that increased resistance reduces the carrying capacity of the population.

The second term in Equation 1 captures cell death from therapy. Drug efficacy is modeled as a Gaussian function [46], where *s* represents the maximum drug efficacy and 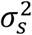 defines the breadth of drug sensitivity. Targeted therapy is modeled as highly specific (*s* = 0.31) and effective over a narrow range of resistance values 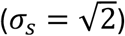, whereas chemotherapy is less specific (*s* = 0.17) but acts over a broader range of traits 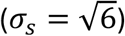. The drug is most effective at *v* = 0, and efficacy decreases as the resistance trait deviates from this value.

The third term in Equation 1 represents the cost of evolvability, modeled as the product of the population’s evolvability (*ε*(*v*)) and a cost coefficient (*d*). This term captures the energetic and regulatory burden of generating heritable variation, such as the cost of increased mutation rates, epigenetic reprogramming, or investment in proteomic and transcriptomic plasticity [47,48]. This cost may also be viewed as the result of producing less fit phenotypes. Alternative ways of modelling this include using partial-differential equations to describe the dynamics of the distribution of phenotypes in the population, thus building in the cost of evolvability (diffusion rate among phenotypes) explicitly [49].

### Constant Evolvability

Next, we define the equations for population and strategy dynamics. The population dynamics (equation 3) is given by the product of the G-function *G*(*v, x*) and the population density (*x*). The rate of evolution of the resistance strategy is constructed in accordance with Fisher’s fundamental theorem of natural selection, according to which the rate of evolution is proportional to the total additive genetic variance of the population, i.e., its evolvability and the selection pressure on the population [6,13,46]. Here, *ε*(*v*) represents the population’s evolvability and 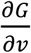 represents the selection gradient.

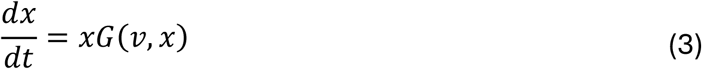

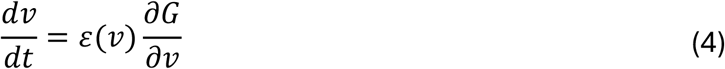

In equation 4, for fixed evolvabilities, we set *ε*(*v*) to a constant (*ε*(*v*) = *ε*). For high evolvability, *ε* = 0.35 and for low evolvability, *ε* = 0.2 When evolvability is facultative, then we consider *ε*(*v*) as a function of *v*.

The condition for eco-evolutionary stability is then defined by: 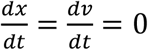 [42,50].

### Facultative Evolvability

Here, we expand the model to consider facultative evolvability. In this case, the evolvability (*ε*(*v*)) of the cancer population changes in response to therapy, as shown in equation 5.

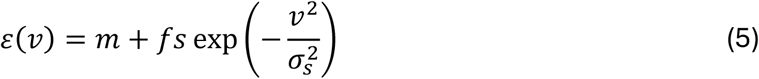

The constant *m* is the baseline mutational burden that accounts for the minimum evolvability in the population [17,51,52]. The second term corresponds to drug-induced death, *s exp* 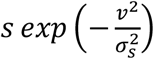, scaled by a factor *f*. We introduce this evolvability scaling factor to ensure that the total evolvability in the facultative population does not exceed that of a population with high constant evolvability (*ε* (*high*) = 0.35). In other words, *f* constrains the maximum evolvability achievable under each therapy type, reflecting biological limits on mutation rates induced by treatment.

So, for choosing *f*, the following condition should hold true-

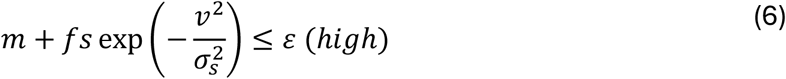

i.e.,

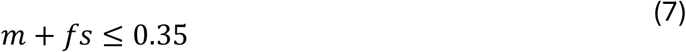

We summarize the variables and parameters used in this model in Tables 1 and 2. The value of *f* will be different under chemotherapy and targeted therapy and can be found in Table 2.

**Table 1:** Summary of variables and symbols used in the single therapy model.

| Symbol | Meaning |
| --- | --- |
| $x$ | Cancer population size |
| $v$ | Resistance strategy |
| $G(v, x)$ | Fitness as a function of population size and resistance strategy |
| $K(v)$ | Carrying capacity as a function of resistance strategy |
| $\varepsilon(v)$ | Population evolvability for resistance strategy |

**Table 2:** Summary of parameters used in the single therapy model.

| Symbol | Meaning | Value |
| --- | --- | --- |
| $\varepsilon$ | Constant evolvability | $\begin{cases} 0.2, & \text{if low evolvability} \\ 0.35, & \text{if, high evolvability} \end{cases}$ |
| $f$ | Evolvability scaling factor | $\begin{cases} 1.91, & \text{if chemotherapy} \\ 1.04, & \text{if, targeted therapy} \end{cases}$<br>(see equation 7) |
| $s$ | Therapy efficacy | $\begin{cases} 0.17, & \text{if chemotherapy} \\ 0.31, & \text{if, targeted therapy} \end{cases}$ |
| $\sigma_s$ | Therapy efficacy breadth | $\begin{cases} \sqrt{6}, & \text{if chemotherapy} \\ \sqrt{2}, & \text{if, targeted therapy} \end{cases}$ |
| $K_m$ | Maximum carrying capacity | 100 |
| $r$ | Intrinsic growth rate | 0.28 |
| $\sigma_k$ | Carrying capacity breadth | 10 |
| $d$ | Cost of evolvability | 0.12 |
| $m$ | Baseline mutational burden | 0.025 |

### Dual Therapy Model

#### Integrating evolution of resistance into multi-agent therapy

In the initial model, we considered administration of only one drug. Next, we model the administration of two drugs. We modify the G-function (equation 1) to account for the two types of therapy, targeted therapy and chemotherapy.

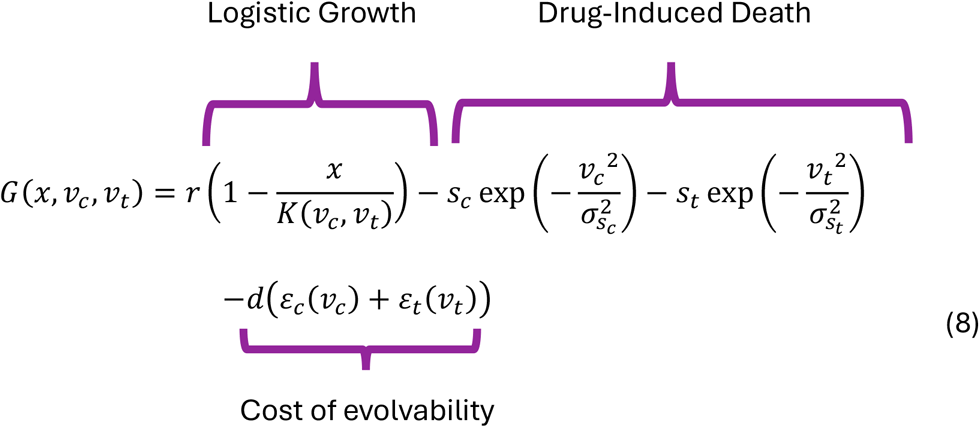

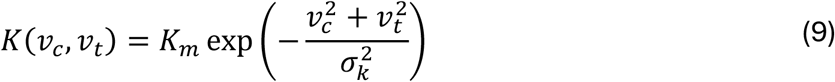

In equation 8, *v*_*c*_ and *v*_*t*_ represent the resistance strategies in response to chemotherapy and targeted therapy, respectively. The first term is the logistic growth term with a modified carrying capacity. The carrying capacity (*K*(*v*_*c*_, *v*_*t*_)) depends on both resistance strategies, ((*v*_*c*_, *v*_*t*_)𝜖 [0, ∞)) (equation 8). *K*(*v*_*c*_, *v*_*t*_) is at its highest (*K*_*m*_) when *v*_*c*_ = *v*_*t*_ = 0. *K*(*v*_*c*_, *v*_*t*_) decreases as the two strategies diverge from 0 in a Gaussian manner.

The second and third terms in equation 8 are the chemotherapy and targeted therapy-induced death terms, respectively. And the last term in equation 8 is the cost of evolvability due to both drugs.

For constant evolvability:

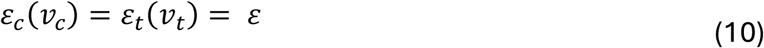

For facultative evolvability:

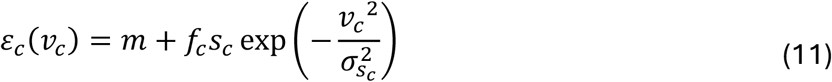

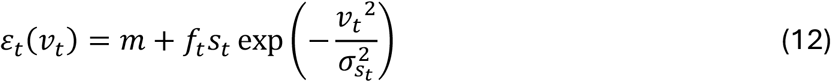

The resistance mechanisms that evolve in response to chemotherapy and targeted therapy could interact in three possible scenarios [46,53]: 1) The two drugs have independent modes of action, and resistance to one drug does not affect resistance to the other. In this case the two drugs are additive. 2) Resistance to one drug confers some resistance to the other drugs, i.e., the two resistance strategies are positively correlated. This leads to cross-resistance. Cross-resistance reduces the efficacy of the two drugs [54,55]. 3) Resistance to one drug makes cells more susceptible to the other drug. The two drugs act synergistically to enhance efficacy beyond additive effects [53,56]. In this case, the two resistance strategies have a negative covariance. Sequential application of such drugs leads to an evolutionary double bind [57]. To model this, we introduce a covariance factor *δ* that captures the inter-dependence of the two resistance strategies [46]. If *δ* = 0 the two resistance strategies evolve independently. If *δ* > 0, the two resistance strategies are positively correlated, and if *δ* < 0, the two strategies are negatively correlated (equations 9 and 10).

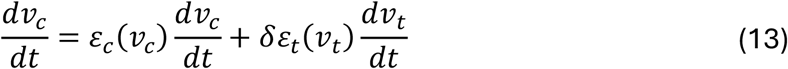

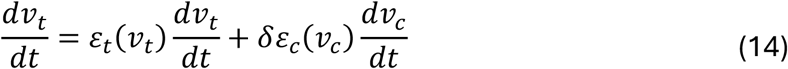

An evolutionary double bind exploits the evolution of resistance by using two drugs where resistance to one increases vulnerability to the other, creating a trap that can drive the cancer population to extinction.

### Combination Therapy

For combination therapy, we use the same model outlined above, but in this case, both drugs are administered simultaneously.

### Simulations

Therapy was administered at time step 600. Before administration of therapy, the resistance strategy does not evolve. The simulations were run until the population and strategy dynamics reached equilibrium. The variable and parameter values for the single-therapy model are summarized in Tables 1 and 2, while those for the dual-therapy model are summarized in Tables 3 and 4.

**Table 3:** Summary of variables introduced in the double bind model.

| Symbol | Meaning |
| --- | --- |
| $v_c$ | Resistance strategy for chemotherapy |
| $v_t$ | Resistance strategy for targeted therapy |
| $G(x, v_c, v_t)$ | Fitness as a function of population size and two different resistance strategies |
| $K(v_c, v_t)$ | Carrying capacity as function of the two resistance strategies |
| $\varepsilon_c(v_c)$ | Evolvability for chemotherapy resistance |
| $\varepsilon_t(v_t)$ | Evolvability for targeted therapy resistance |

**Table 4:** Summary of parameters used in the double bind model.

| Symbol | Meaning | Value |
| --- | --- | --- |
| $s_c$ | Therapy efficacy of chemotherapy | 0.17 |
| $s_t$ | Therapy efficacy of targeted therapy | 0.31 |
| $\sigma_{s_t}$ | Targeted therapy efficacy breadth | $\sqrt{2}$ |
| $\sigma_{s_c}$ | Chemotherapy efficacy breadth | $\sqrt{6}$ |
| $f_c$ | Evolvability scaling factor for chemotherapy | 1.91<br>(see equation 7) |
| $f_t$ | Evolvability scaling factor for targeted therapy | 1.04<br>(see equation 7) |
| $\delta$ | Covariance between two resistance strategies | 0.015 |

## Results

### Adaptive Landscape

We begin our analysis by simulating how chemotherapy and targeted therapy shape the cancer population’s adaptive landscape (Fig. 2). Targeted therapy (red curve) produces a steep drop at *v* = 0, indicating its high specificity against cells with low resistance.

**Figure 2:**
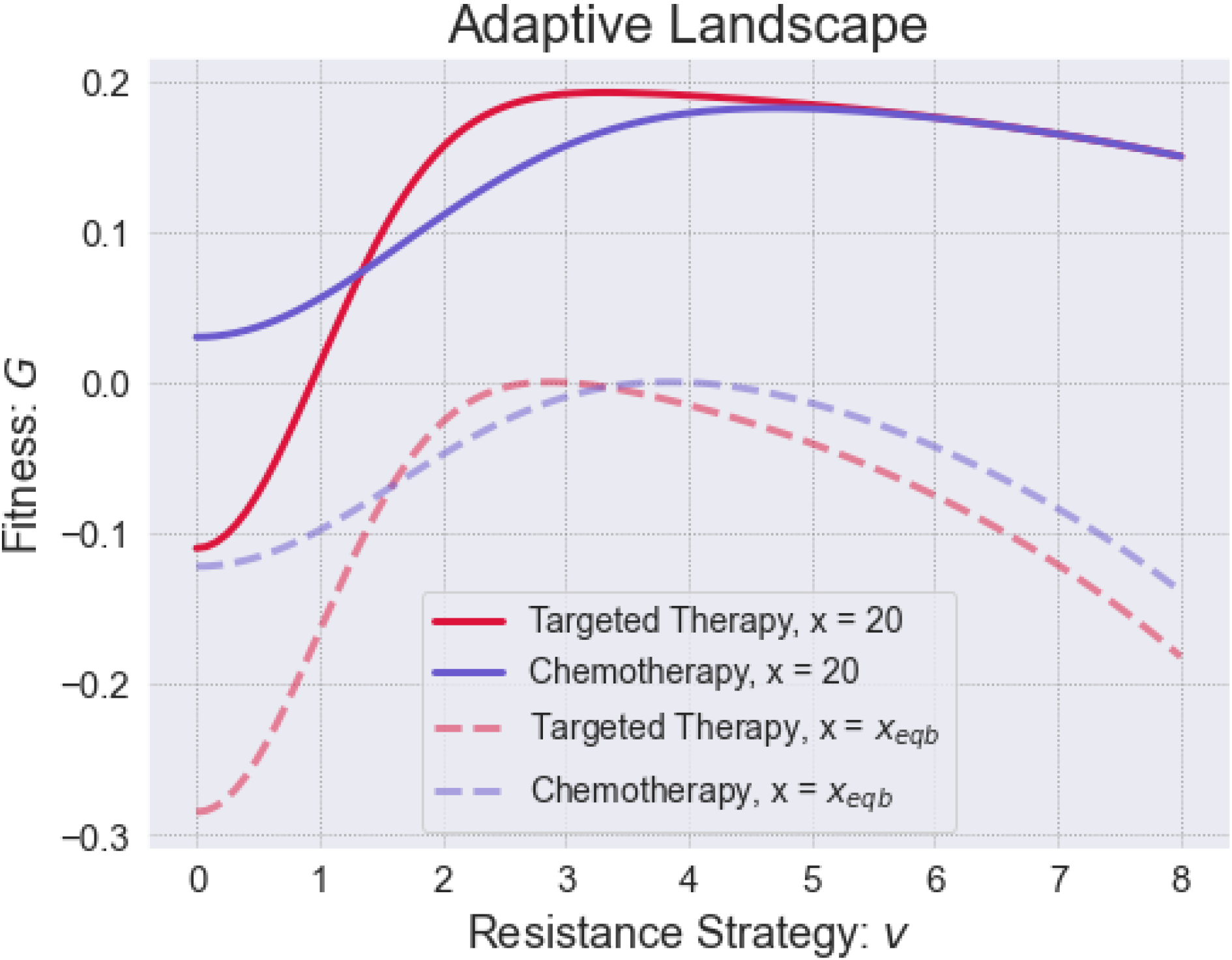
Adaptive Landscape of Cancer Population under Therapy: Here we show the adaptive landscape of the cancer population with fixed low evolvability, under chemotherapy (blue) and targeted therapy (red). Targeted therapy induces a steep selection gradient at strategies close to *v* = 0 and rapid evolution of resistance as the strategy values deviate from *v* = 0. Chemotherapy induces a gentler gradient but is effective on a broader range of resistance strategies. The adaptive landscape is drawn for a fixed population size. The solid lines correspond to, *x* = 20 and the dashed lines correspond to the adaptive landscape at equilibrium. For chemotherapy, *x*_*eqb*_ = 74.32 and for targeted therapy *x*_*eqb*_ = 82.47.

The negative fitness at *v* = 0 indicates that the population will go extinct under targeted therapy unless resistance evolves. Its efficacy rapidly decreases as the resistance values diverge from 0. This steep gradient in the landscape represents a strong selection pressure, which can accelerate the evolution of resistance. The rapid evolution of resistance would render therapy ineffective through evolutionary rescue.

In contrast, chemotherapy (blue curve) results in a shallower drop in the adaptive landscape but leads to lower fitness for a bigger range of resistance values. This is consistent with its broad-spectrum activity against a larger range of cancer phenotypes. The gentler selection gradient means that resistance to chemotherapy evolves more slowly. To become resistant to chemotherapy, cancer cells must adopt more extreme values of resistance that are costlier and reduce overall fitness. The adaptive landscape for targeted therapy (at *x* = 20) reaches a peak at *v* = 3.13. For chemotherapy, the peak is at *v* = 4.68. Beyond this peak, fitness decreases again due to the increasing cost of resistance.

Based on these observations, we hypothesize that for cancer populations with low evolvability (slow-evolving cancers), targeted therapy may be more effective due to its high efficacy at low resistance levels. Populations with high evolvability (fast-evolving cancers) can rapidly evolve resistance to targeted therapy. In contrast, chemotherapy would drive them toward more extreme and costly resistance strategies, which would reduce their overall fitness. In the following sections, we will explore these therapy regimens.

### Constant Evolvability

We simulate the administration of targeted therapy and chemotherapy in Fig. 3. In the absence of therapy, low-evolvability cancers maintain a higher tumor burden than the high-evolvability one. This is due to the greater cost of evolvability, which reduces the effective carrying capacity of the high-evolvability population (85.05 vs 91.46, see Fig. 3a & b).

**Figure 3:**
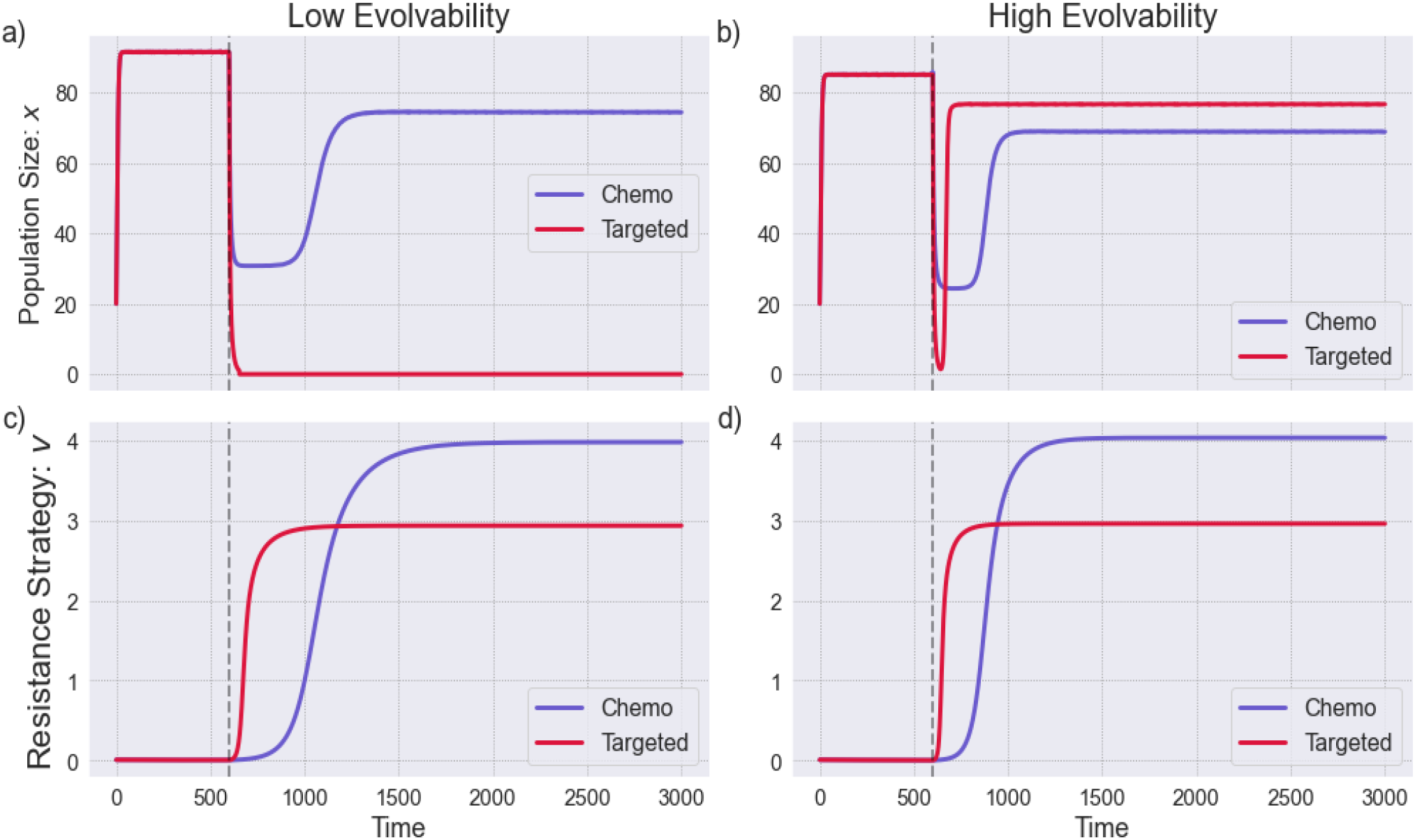
Dynamics of cancer populations with fixed evolvabilities: Population dynamics (a,b) and resistance strategy dynamics (c,d) for the cancer population with low constant evolvability (a,c) and high constant evolvability (b,d). Therapy is administered at t = 600 (shown by the dashed vertical line). If the population size dropped below 1 at any point, we imposed an extinction threshold, setting the population to zero for all subsequent time steps. Without this extinction threshold, the population size at equilibrium for targeted therapy would be 82.47.

Once therapy is administered, the slow-evolving population is unable to adapt rapidly to targeted therapy and is driven to extinction. Had it been able to avoid extinction, it would have evolved up its adaptive landscape to an eco-evolutionary equilibrium (red dashed line in Fig. 2). In contrast, the fast-evolving population undergoes evolutionary rescue and achieves an eco-evolutionary equilibrium population size of *x* = 76.59 and resistance strategy of *v* = 3.87 (blue dashed line in Fig. 2). Under chemotherapy, both populations evolve higher resistance strategies (Fig. 3c and 3d). The period of remission is longer for the slow evolver than the fast evolver. However, the fast-evolving population reaches greater resistance values than the slow-evolving one, resulting in a higher cost of resistance. This, combined with the higher intrinsic cost of evolvability, leads to a greater reduction in overall fitness. As a result, the fast-evolving population stabilizes at a lower equilibrium size (68.79) compared to the slow-evolving population (74.32).

These findings indicate that targeted therapy drives the slow-evolving population to extinction due to negative fitness under therapy and the lack of a sizable population and time to evolve quickly enough to a resistance strategy with positive fitness. Chemotherapy, on the other hand, is more effective at suppressing the fast-evolving population by forcing it to adopt costly resistance strategies.

### Facultative Evolvability

We now compare the ecological and evolutionary trajectories of cancer populations with constant evolvability to those with facultative evolvability (Fig. 3). Facultative evolvability allows a population to dynamically adjust its capacity to generate heritable variation in response to environmental conditions. In our model, environmental stress is induced by the administration of therapy. In response, cancer cells may activate mechanisms that increase their evolvability [58,59]. While increased evolvability can enhance a population’s ability to acquire resistance, it also imposes a greater cost of evolvability and resistance. The facultative population must therefore balance the trade-off between achieving high resistance and minimizing the associated metabolic and fitness costs [60].

Prior to drug administration, the facultative population maintains its evolvability at a baseline mutation rate. By staying at this baseline level, the population minimizes the cost of evolvability. As a result, the facultative population achieves a higher effective carrying capacity (98.9) compared to populations with fixed evolvability, which continuously pay a higher cost regardless of the presence or absence of therapy.

After the administration of therapy, the evolvability of the facultative population increases dramatically (Fig. 4c and 4f). This surge enables a rapid increase in the resistance strategy, allowing the population to adapt quickly. Resistance evolves more rapidly in response to targeted therapy than to chemotherapy, consistent with the predictions from the adaptive landscapes in Fig. 2. Once a sufficient level of resistance is achieved, evolvability begins to decline. This decline occurs more rapidly under targeted therapy than chemotherapy, reflecting the faster adaptation dynamics in the former.

**Figure 4:**
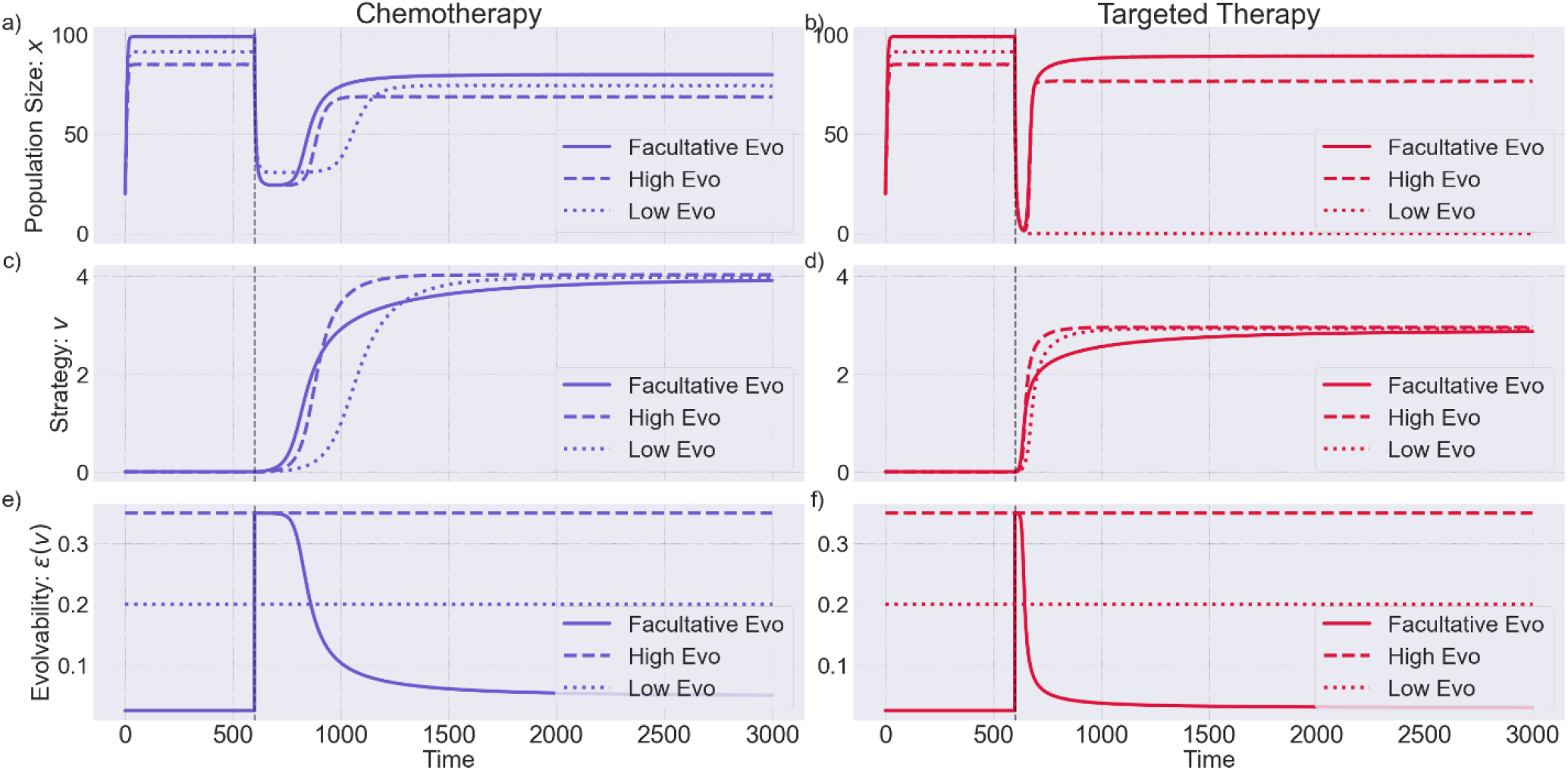
Comparison of dynamics with fixed and facultative evolvability: Population dynamics (a,b), resistance strategy dynamics (c,d), and evolvability (e,f) dynamics of the slow-evolving, fast-evolving, and facultative cancer population. Therapy is administered at t = 600 (indicated by the vertical dashed line).

Our simulations show the facultative population attains a higher equilibrium population size than the populations with fixed evolvability (79.86 in the case of chemotherapy and 89.23 in the case of targeted therapy). Facultative evolvability would increase the chances of therapy failure and relapses in patients.

### Intermittent Therapy

To further explore the therapeutic implications of evolvability, we simulate intermittent therapy, where treatment is administered in periodic cycles rather than continuously. Intermittent therapy has been investigated clinically as a strategy to reduce toxicity, delay resistance, and prolong treatment efficacy [61,62]. By periodically withdrawing therapy, this approach reduces the selective pressure that drives the emergence of resistant clones. In our model, the cyclic nature of intermittent therapy creates alternating periods of stress and recovery, and we examine how populations with fixed versus facultative evolvability adapt under this dynamic treatment regimen.

Figs. 5 and S1 show simulations for intermittent targeted therapy and chemotherapy, respectively, on the slow-evolving (low fixed evolvability), fast-evolving (high fixed evolvability), and the facultatively-evolving population. Therapy is administered at the 600th time-step and removed and restarted every 50 time steps. During the ‘on therapy’ periods, the population size decreases and resistance increases. During the ‘off therapy’ or ‘drug holiday’ periods, the cancer population bounces back, and the level of resistance decreases. This regular cycling between treatment and rest leads to fluctuations in both population size and resistance strategy dynamics. The qualitative results under intermittent therapy overall match those under continuous therapy. For populations with fixed evolvability, targeted therapy achieves near-curative response. However, once therapy is revoked after 50 time steps, the population recovers and stabilizes into oscillations between a size ∼ 81–87. A longer cycle duration (or continuous therapy as in Fig. 3) would drive this population extinct. This highlights the importance of timing the drug holidays right.

**Figure 5:**
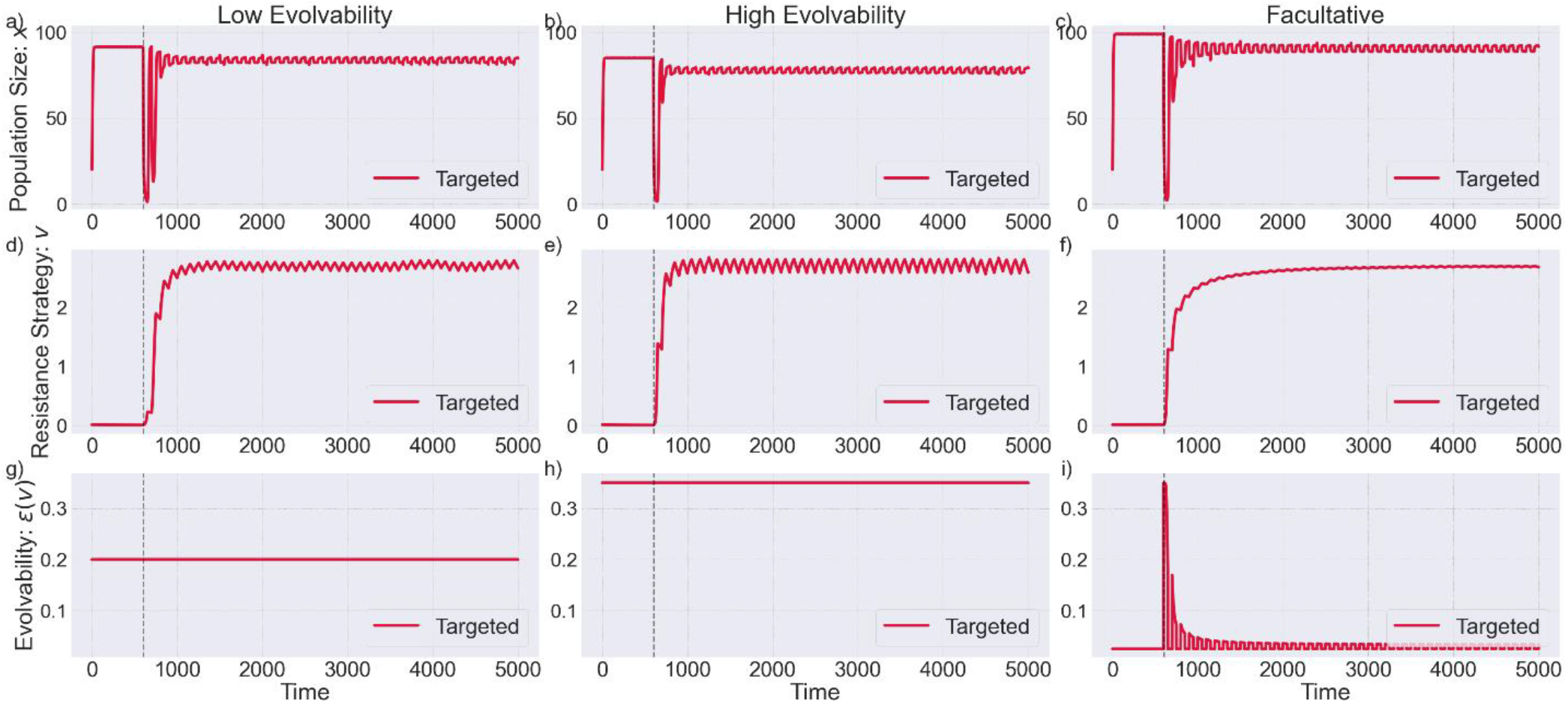
Comparison of dynamics under intermittent targeted therapy: Population dynamics (a,b,c), resistance strategy dynamics (d,e,f), and evolvability (g,h,i) dynamics of the slow-evolving, fast-evolving, and facultative cancer population under intermittent therapy. Therapy is administered at t = 600 (indicated by the vertical dashed line) and turned ‘on’ or ‘off’ every 50 time-steps

The fast-evolving population oscillates between a population size of ∼76-79 under targeted therapy. However, unlike the slow-evolving population, extending cycle duration does not drive this population to extinction (as seen in the continuous therapy case in Fig. 3). Increasing therapy efficacy is the only way to eliminate it using a single drug. Note that with high evolvability, the proportional change in the cancer cells’ resistance strategy exceeds changes in their population size.

With chemotherapy (Fig. S1), natural selection drives both slow evolvers and fast evolvers to adopt more extreme resistance strategies than under targeted therapy. The fast-evolving population evolves higher resistance levels than the slow-evolving one, incurring a greater fitness cost. As a result, it oscillates between ∼68-76 at equilibrium, while the slow-evolving population cycles between ∼74-82. The smaller amplitude of oscillation and lower overall tumor burden in the fast-evolving population suggest better tumor control under chemotherapy. As expected, the slow evolver’s resistance strategy cycles with a lower amplitude than the fast evolvers.

Consistent with our earlier observations in Fig. 3, the facultative population fares better in response to therapy than populations with fixed evolvability. Under targeted therapy, it reaches an equilibrium population size ranging from ∼88-92, while under chemotherapy, the range is slightly lower, between ∼79-87. In both cases, the facultative population achieves higher equilibrium sizes than its fixed-evolvability counterparts. Before drug administration, the facultative cells maintain evolvability at a low baseline level to minimize fitness costs. Following therapy administration, evolvability spikes sharply, enabling rapid adaptation, and then decreases again during drug holiday periods.

Curiously, for the facultative evolver, the population’s resistance strategy remains nearly constant with much smaller fluctuations than either of the fixed evolvers. At first, this seems counterintuitive, as a benefit of facultative evolvability would be to adjust the resistance strategy quickly to conform to the therapy circumstances. Yet, evolving rapidly to the on-off cycles of therapy incurs a high cost of evolvability. With on-off cycles of 50, the facultative evolvers achieve the highest fitness by adopting a nearly fixed resistance strategy and thus minimizing the cost of evolvability. The fixed evolvers do not have this opportunity to balance evolvability with the cost of evolvability, as they pay a constant cost of evolvability regardless of the circumstances.

### Double Bind

Our next goal is to design a therapy regimen capable of driving the high evolvability and the facultative evolvability population to extinction. To this end, we explore an evolutionary double bind — a strategy that traps the cancer population with no viable path to escape, much like a mouse caught between an owl in the open field and a snake in the bushes [63]. This approach involves the sequential, cyclic administration of two drugs, such that resistance to one drug increases sensitivity to the other. Paramount to the design of an evolutionary double bind is the choice of drug order and cycle duration. The sequence in which drugs are administered may influence the rate at which resistance evolves and treatment efficacy. Equally critical is the timing of switching therapies. For example, the first drug may drive the population into an evolutionary bottleneck; if the second drug is administered before resistant clones emerge, it can eliminate the remaining vulnerable cells and drive the population to extinction. This narrow window, before evolutionary rescue occurs, is the cure window. Drug timing is key. If the cycle is too short, resistance may not have evolved sufficiently to trigger the double bind; if too long, the population may adapt and escape, making cure impossible.

The potential for double binds have been explored by combining chemotherapy with immunotherapy, and radiotherapy with immunotherapy [64,65]. In this section, we simulate an evolutionary double bind using chemotherapy and targeted therapy to investigate how the order of drug administration and cycle duration influence the evolutionary dynamics of the cancer population. We reason that administering chemotherapy first may slow down the evolution of resistance, as suggested by the shallower selection gradient observed in the adaptive landscape (Fig. 2). If targeted therapy is administered after chemotherapy, it could eliminate cancer cells before they have time to adapt. We test this strategy in Fig. 6 using a cycle duration of 50 time steps.

**Figure 6:**
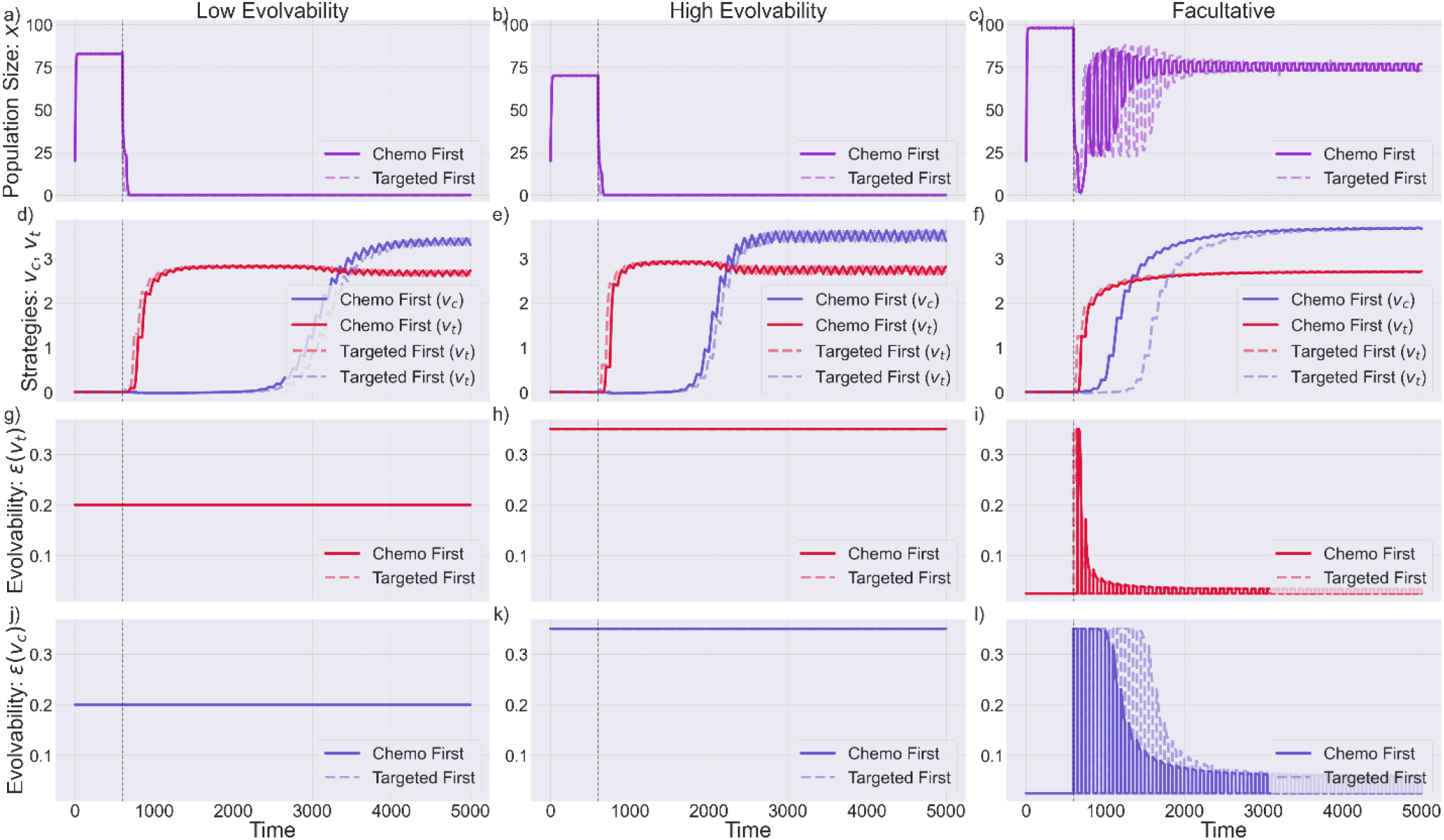
Comparison of dynamics under double bind therapy: Population dynamics (a,b,c), resistance strategy dynamics (d,e,f), and evolvability (g-l) dynamics of the slow-evolving, fast-evolving, and facultative cancer population under double bind therapy. Therapy is initiated at t=600 (indicated by the vertical dashed line). Solid lines represent the regimen where chemotherapy is given first, followed by targeted therapy; dashed lines represent the reverse order. The cycle duration is short, i.e., therapy is switched every 50 steps.

First, note that prior to the administration of therapy, the effective carrying capacity of all three populations is lower in the double bind case compared to the continuous (single therapy) cases discussed above. This reduction arises because the double bind model includes two resistance strategies and two evolvabilities, which imposes a higher cumulative cost on the population. The drop in effective carrying capacity is most pronounced in the high-evolvability population (69.9 in the double bind vs. 91.4 in the continuous case), followed by the low-evolvability population (82.8 vs. 85.05), and finally the facultative population (97.8 vs. 98.9). Interestingly, even before therapy begins, the facultative population performs better than the fixed-evolvability populations. This is because it maintains evolvability at a baseline level, avoiding the higher costs incurred by constantly sustaining high evolvability.

Using a double bind of cycle duration 50 (short cycle), both the low and high fixed evolvability populations are driven to extinction. Chemotherapy slows down the evolution of resistance and brings the population down. At this point, when targeted therapy is administered, with its higher efficacy, it drives the population to 0, i.e., cure. However, this strategy fails in the case of facultative evolvability.

In the facultative case, evolvability itself fluctuates cyclically. When chemotherapy is administered first, *ε*_*c*_(*v*_*c*_) (evolvability for chemotherapy resistance) increases, while *ε*_*t*_(*v*_*t*_) (evolvability for targeted therapy resistance) remains at its baseline level. Upon switching to targeted therapy, *ε*_*t*_(*v*_*t*_) increases and *ε*_*c*_(*v*_*c*_) decreases back to baseline. This cycling continues every time therapy is switched, and the amplitude of oscillations gradually dampens as the population acquires sufficient resistance to ensure survival. Because of this rapid and alternating modulation of evolvability, the cross-susceptibility between resistance strategies is not fully exploited, rendering the double bind ineffective.

While resistance to therapy is undesirable, in a double bind, the emergence of just the right amount of resistance is necessary to make the population susceptible to the sequential application of drugs. A short cycle length fails to induce a complete resistance response and, as a result, does not effectively set the evolutionary trap. To allow enough time for resistance to evolve and for one type of therapy to counteract the efficacy of the other, we increased the cycle duration to 200 (Fig. 7). A longer cycle duration allows enough time for resistance to the first drug to develop, creating a vulnerability that the second drug can exploit, even in the facultative population. We see that with this cycle duration, the facultative population is driven to extinction in the second cycle when targeted therapy is applied following chemotherapy. Reversing the order in which the drugs are applied enables evolutionary rescue.

**Figure 7:**
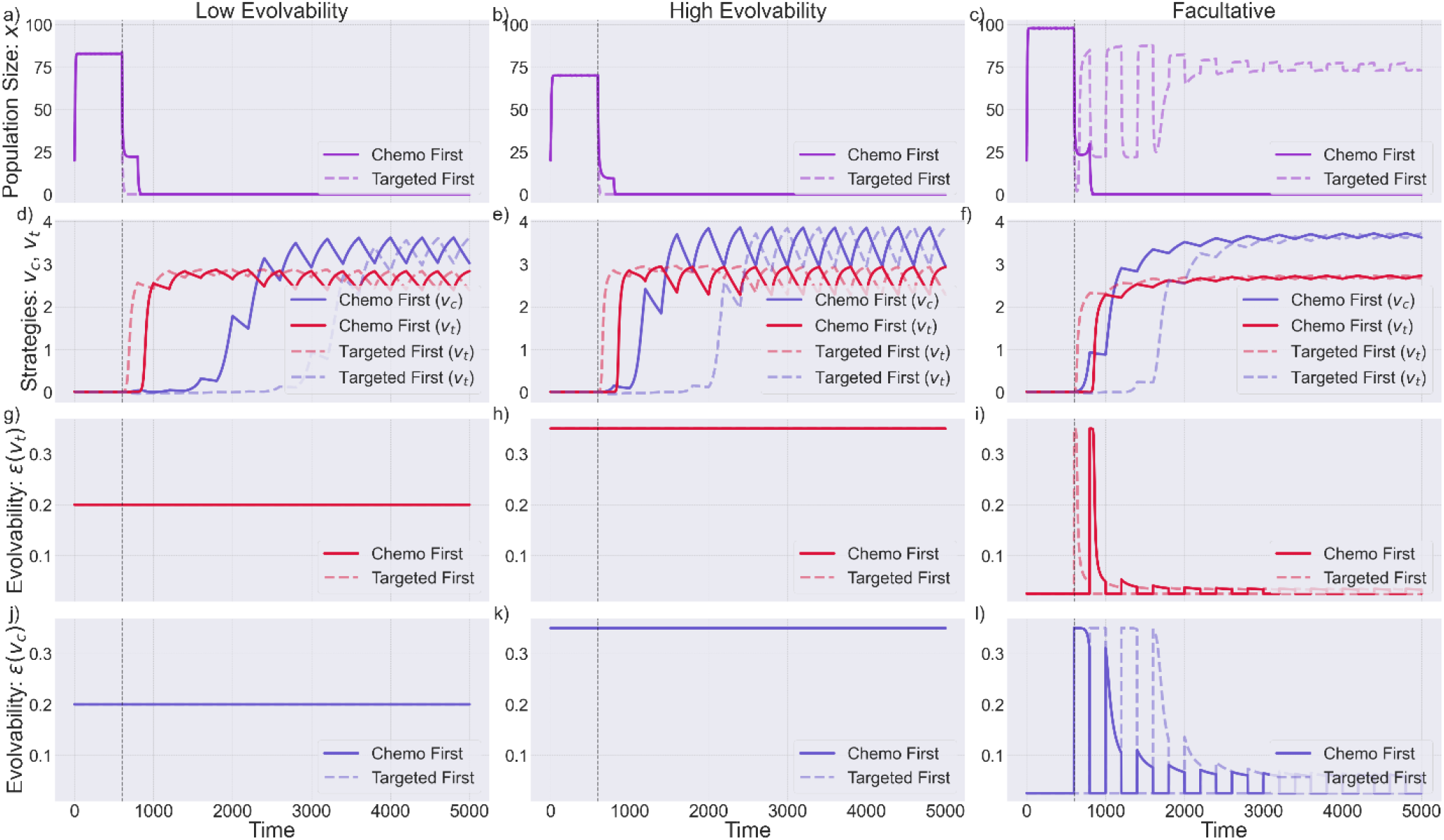
Comparison of dynamics under double bind therapy: Population dynamics (a,b,c), resistance strategy dynamics (d,e,f), and evolvability (g-l) dynamics of the slow-evolving, fast-evolving, and facultative cancer population under double bind therapy. Therapy is initiated at t=600. Solid lines represent the regimen where chemotherapy is given first, followed by targeted therapy; dashed lines represent the reverse order. The cycle duration is long, i.e., therapy is switched every 200 steps.

To facilitate comparison, we present results for combination therapy, where chemotherapy and targeted therapy are administered simultaneously, in Fig. S2. This approach, continuous administration of a drug cocktail without sequencing or switching, is commonly used in clinical settings. When the full drug dose is applied, all three populations are driven extinct. However, in practice, drug doses are often reduced to avoid excessive toxicity. To reflect this, we also simulate treatment dynamics at half the full dose. Under these conditions, the high-evolvability population is still eliminated, but the low-evolvability and facultative-evolvability populations escape cure. While in theory, combination therapy with full doses is the best therapy option, sequential double bind therapy is superior to half-dose combination therapy and prevents toxicity associated with administering very high drug doses together.

We provide another example of the design of double bind therapy in the supplementary section. This cancer population has a higher intrinsic growth rate and a lower cost of evolvability. We set the evolvability value (*ε*) to 0.15 for the low fixed-evolvability population and increase it to 0.4 for the high fixed-evolvability population (see Table S1). In this case, with a short cycle duration of 50, both the low-evolvability and facultative-evolvability populations undergo evolutionary rescue and escape the double bind trap (Fig. S3). To test whether a longer cycle would improve outcomes, we increase the cycle duration to 200, like in the previous case. Under this regimen, the facultative population is eliminated, but the low-evolvability population still escapes extinction. This presents an interesting case where low fixed evolvability offers a survival advantage over facultative evolvability under double bind therapy. The reason lies in the lower cost of evolvability for the low fixed-evolvability population. The facultative population rapidly increases its evolvability in response to therapy, which temporarily reduces its fitness and makes it more vulnerable to the trap (Fig. S4).

The most straightforward way to eliminate the low-evolvability population is to either increase therapy efficacy or use combination therapy (Fig. S5). However, we also explore whether cure can be achieved using a double bind strategy alone. By extending the cycle duration to 500, we are able to drive the low-evolvability population to extinction (Fig. S6). These results suggest that to eliminate this population, one must either increase drug efficacy or prolong the treatment cycle. Depending on clinical constraints like toxicity and how quickly cure is desired, a choice can be made between stronger or longer treatment.

Overall, cancer evolvability plays a critical role in shaping the response to therapy, and treatment strategies should be designed accordingly. By understanding the evolutionary dynamics of resistance and the influence of evolvability, we can make informed decisions about whether to combine therapies or sequence them. When sequencing drugs in a double bind strategy, allowing just the right amount of resistance to evolve is essential for setting the trap. This process is governed by the drug order, timing of the switch, and cycle duration—all of which must be carefully optimized to maximize treatment efficacy.

## Discussion

In this study, we explored how cancer evolvability shapes therapeutic outcomes using an evolutionary game theory framework. Evolvability-the heritable variation a population can generate-serves as the fuel for natural selection [7,14]. In cancer, such variation arises from genetic mutations, epigenetic modifications, and non-genetic mechanisms such as transcriptional noise and cell-state plasticity [25,66]. These processes enable tumor cells to adapt to environmental changes, including therapeutic pressure.

We specifically examined how resistance to chemotherapy and targeted therapy evolve. Targeted therapies act on specific oncogenic pathways [34]. They impose steep selection gradients that accelerate the evolution of resistance. In contrast, chemotherapy targets broad cellular processes like DNA replication and mitosis [29]. This results in a flatter adaptive landscape, with resistance evolving slowly but reaching higher levels. These dynamics align with clinical observations: while resistance to chemotherapy tends to emerge more slowly, it is often more aggressive and confers cross-resistance to multiple agents [54]. Our results demonstrate that targeted therapies can eliminate slow-evolving tumors with low, fixed evolvability. However, tumors with high evolvability often survive by undergoing evolutionary rescue. In such cases, chemotherapy proves more effective by selecting for extreme resistance strategies that carry higher fitness costs.

We also examined therapy outcomes for cancers with facultative evolvability where, a cell’s capacity to generate variation depends on therapy-induced stress [38]. Cancer cells, for instance, can accrue more mutations and phenotypic plasticity in response to therapy [24,58]. This increase in the ability to generate variation can help the population adapt faster and better to a dynamically changing environment. Our simulations show that, by dynamically balancing the trade-off between resistance and the associated costs of resistance and evolvability, facultative cells generally attain higher fitness than their fixed-evolvability counterparts [60].

To target the adaptability of facultative cells, we implement an evolutionary double bind — a strategy that exploits the negative covariance between resistance mechanisms, such that resistance to one drug increases sensitivity to the other [55–57]. We find that, with the right order and cycle duration of drug administration, the facultative population can be driven to extinction. Administering chemotherapy first slows down the rate of evolution of resistance. Following this with the administration of targeted therapy drives the facultative population to extinction. The reverse order fails to achieve the same outcome.

Basanta *et al*. had similar findings to ours [65]. Using a matrix game–based model to simulate an evolutionary double bind involving chemotherapy and a *p53* vaccine, they showed that administering chemotherapy first, followed by the vaccine, reduced tumor heterogeneity and improved therapeutic outcomes [43,67]. More recently, using a Lotka-Volterra competition model [66], Luddy *et al*. [64] demonstrated that radiation therapy followed by NK cell–based immunotherapy creates a double bind. Radiation selects for cells expressing NK cell ligands, which are then effectively targeted by immunotherapy. Their model was supported by experimental validation, where combination therapy was less effective than sequential therapy. Both studies highlight the critical role of drug order, timing, and duration in designing effective evolutionary therapies.

Our study uses a continuous game framework where resistance is a continuous trait rather than binary (sensitive vs. resistant) [42]. Our approach is similar in spirit to the work of Cunningham *et al*. [63], who incorporated intrinsic and environmental resistance, as well as phenotypic plasticity, to examine the efficacy of double bind therapy. We specifically investigate the role of evolvability in shaping therapeutic response and guiding treatment choice. Unlike previous models, ours does not rely on explicit drug sensitization parameters. Instead, we introduce a covariance term that captures how resistance to one drug influences the rate of adaptation to another [46].

In summary, there is growing recognition of the role that evolutionary processes play in driving cancer progression and therapeutic resistance. A key focus has been on quantifying the rate of clonal evolution shaped by genetic mutations, epigenetic remodeling, chromosomal instability, and non-genetic phenotypic plasticity, all of which contribute to a tumor’s evolvability [20,24,28,59,69–71]. Different cellular traits may exhibit different evolvabilities, meaning that resistance mechanisms to the same therapy, even within the same cell, can evolve at distinct rates. Recent advances in molecular biology and high-throughput technologies have enabled more precise characterization of these mechanisms [17]. For example, single-cell RNA sequencing (scRNA-seq) and deep mutational scanning allow for high-resolution mapping of mutational landscapes and evolutionary trajectories within heterogeneous tumors [72,73]. Live-cell imaging and multiplexed spatial profiling can reveal dynamic phenotypic heterogeneity and cell-state transitions over time [69,74–76]. Additionally, techniques such as ChIP-seq and ATAC-seq provide insights into epigenetic modifications and chromatin accessibility that regulate gene expression variability and potential for adaptation [77–79]. Together, these tools are beginning to make it possible to quantify evolvability as a biomarker and incorporate it into therapeutic decision-making.

An additional and critical feature of the Darwinian dynamics underlying clinical resistance is the ability of cancer cells to modulate their evolvability in response to stress, i.e., facultative evolvability. This dynamic adaptability is an additional layer of complexity to therapeutic outcomes and underscores the limitations of conventional resistance models that assume fixed traits. Experimental advances now make it possible to quantify the biological mechanisms that contribute to evolvability. These measurements can be integrated with mathematical models to simulate resistance dynamics, optimize treatment schedules, and inform therapeutic design. For strategies like evolutionary double binds, success depends not only on drug choice but also on the order and timing of administration [57,80]. As precision oncology continues to advance, incorporating assessments of tumor evolvability into clinical decision-making may become essential. While more work is needed to translate these insights into practice, evolution-informed therapy design offers a promising path toward delaying resistance and improving patient outcomes.

## Acknowledgements

The authors thank Chris Whelan for proofreading and providing suggestions that improved the manuscript. R.B. also acknowledges Praneet Srivastava for insightful discussions, brainstorming, and troubleshooting ideas during the project.

The authors used Grammarly and ChatGPT for copy-editing the manuscript. The authors acknowledge Moffitt Cancer Center’s academic institution license for BioRender.com.

## Data Availability

Codes used to generate simulations in the paper are available at https://github.com/rbhattacharya49/Evolvability-Determines-Therapy-Outcomes

## Author Contributions

A.B. conceptualized the study. R.B. wrote the manuscript and created the figures. R.B., A.B., J.B., and R.G. reviewed and edited the manuscript.

