## Supplementary Information and Figures for "Cancer Evolvability Determines Therapy Outcomes"

### Supplementary Section

Ranjini Bhattacharya[1,2*], Anuraag Bukkuri [3], Robert A. Gatenby [4,5], Joel S. Brown[1,5]

1 Department of Integrated Mathematical Oncology, H. Lee Moffitt Cancer Center and Research Institute, Tampa, FL, USA

2 Department of Cancer Biology, University of South Florida, Tampa, FL, USA

3 Department of Mathematics, City St. George's, University of London, London, UK

4 Department of Radiology, H. Lee Moffitt Cancer Center and Research Institute, Tampa, FL, USA

5 Cancer Biology and Evolution Program, Moffitt Cancer Center and Research Institute, Tampa, FL, USA

Intermittent Therapy

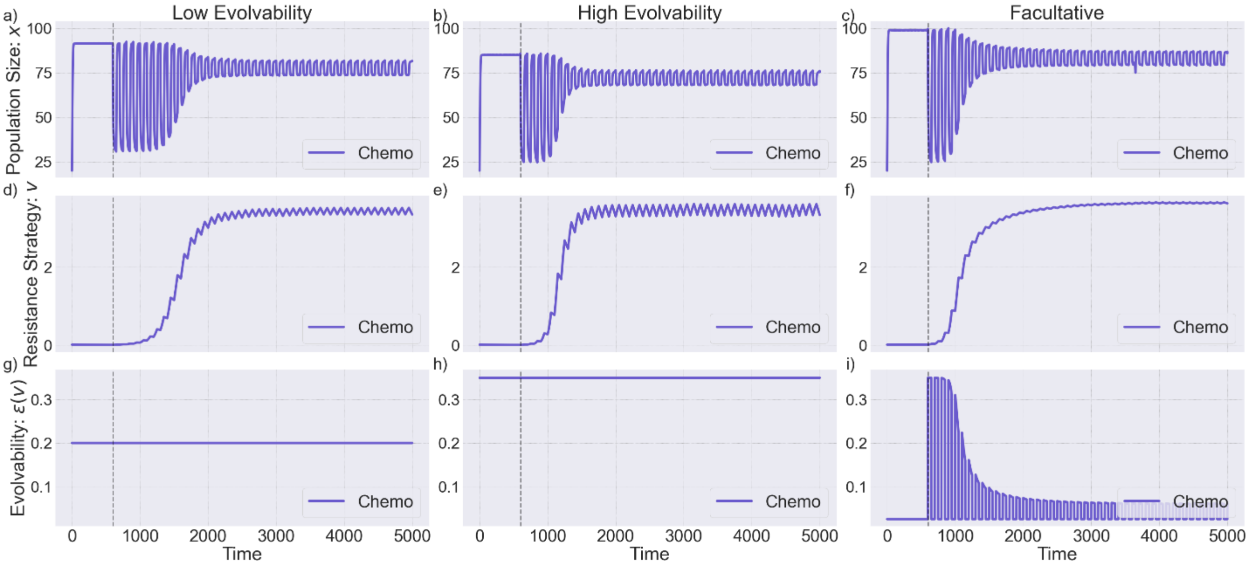


Figure S1: Comparison of dynamics under chemotherapy: Population dynamics (a,b,c), resistance strategy dynamics (d,e,f), and evolvability (g,h,i) dynamics of the slow-evolving, fast-evolving, and facultative cancer population under intermittent therapy. Therapy is administered at t = 600 and turned 'on' or 'off' every 50 time-steps. Parameters used are shown in Table 1.

Combination Therapy

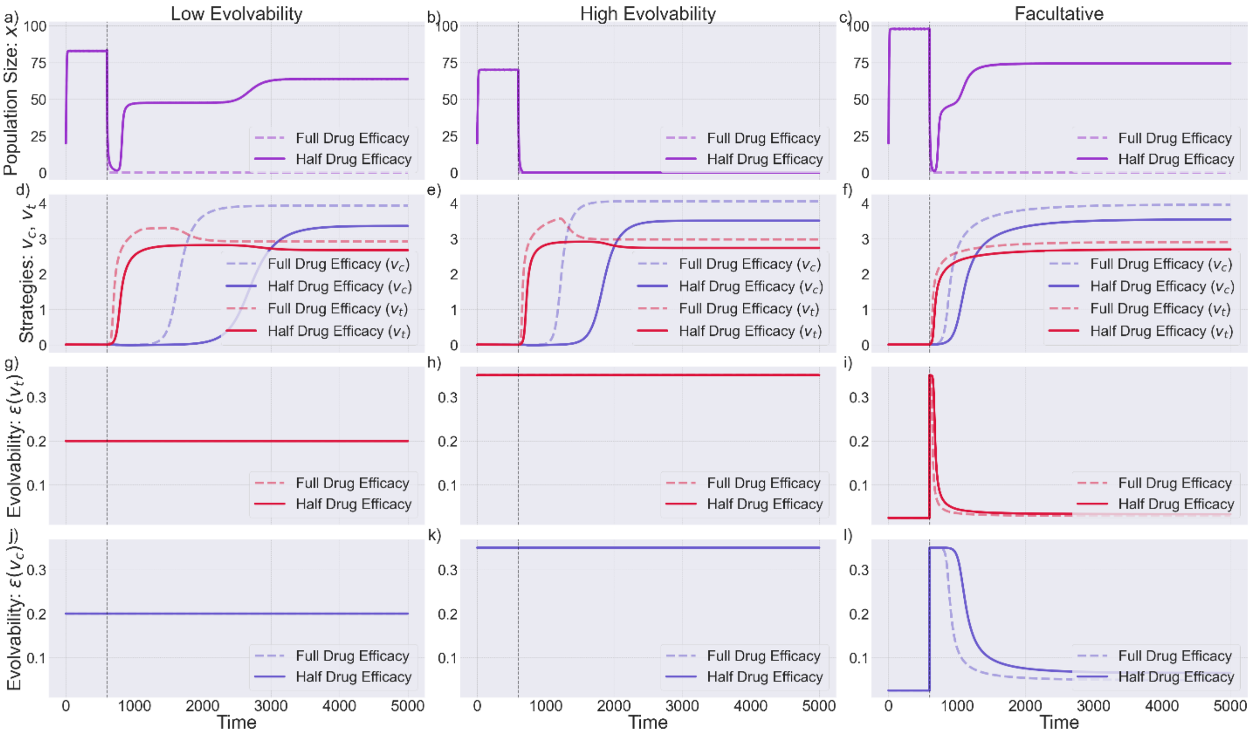


Figure S2: Comparison of dynamics under combination therapy: Population dynamics (a,b,c), resistance strategy dynamics (d,e,f), and evolvability (g-l) dynamics of the slow-evolving, fast-evolving, and facultative cancer population under combination therapy. Therapy is initiated at $t = 600$. Solid lines represent the regimen where the full drug dose is administered; dashed lines show the regimen where half of the drug dose is administered to reduce toxicity. Parameters used are shown in Table 1 & 2.

Alternative Case:

Table S1 shows the parameters used for Fig. S2 and S3.

| Symbol | Meaning | Value |
| --- | --- | --- |
| $r$ | Intrinsic growth rate | **0.32** |
| $s_c$ | Therapy efficacy of chemotherapy | 0.17 |
| $s_t$ | Therapy efficacy of targeted therapy | **0.3** |
| $\sigma_{s_t}$ | Targeted therapy efficacy breadth | $\sqrt{2}$ |
| $\sigma_{s_c}$ | Chemotherapy efficacy breadth | $\sqrt{6}$ |
| $f_c$ | Evolvability scaling factor for chemotherapy | **2.20** |
| $f_t$ | Evolvability scaling factor for targeted therapy | 1.25 |
| $\delta$ | Covariance between two resistance strategies | 0.015 |
| $d$ | Cost of evolvability | **0.09** |
| $m$ | Baseline mutational burden | 0.025 |
| $K_m$ | Maximum carrying capacity | 100 |
| $\sigma_k$ | Carrying capacity breadth | 10 |
| $\varepsilon$ | Constant evolvability | $\begin{cases} \mathbf{0.15,\ \mathit{if}\ low\ evolvability} \\ \mathbf{0.4,\ \mathit{if},high\ evolvability} \end{cases}$ |
| $\delta$ | Covariance between two resistance strategies | 0.015 |

Table S1: Summary of parameters used in the alternative double bind simulations. Parameters that have changed from the original model (see table 1 & 2) are highlighted in bold.

Alternative Case: Double Bind

Cycle Duration: 50

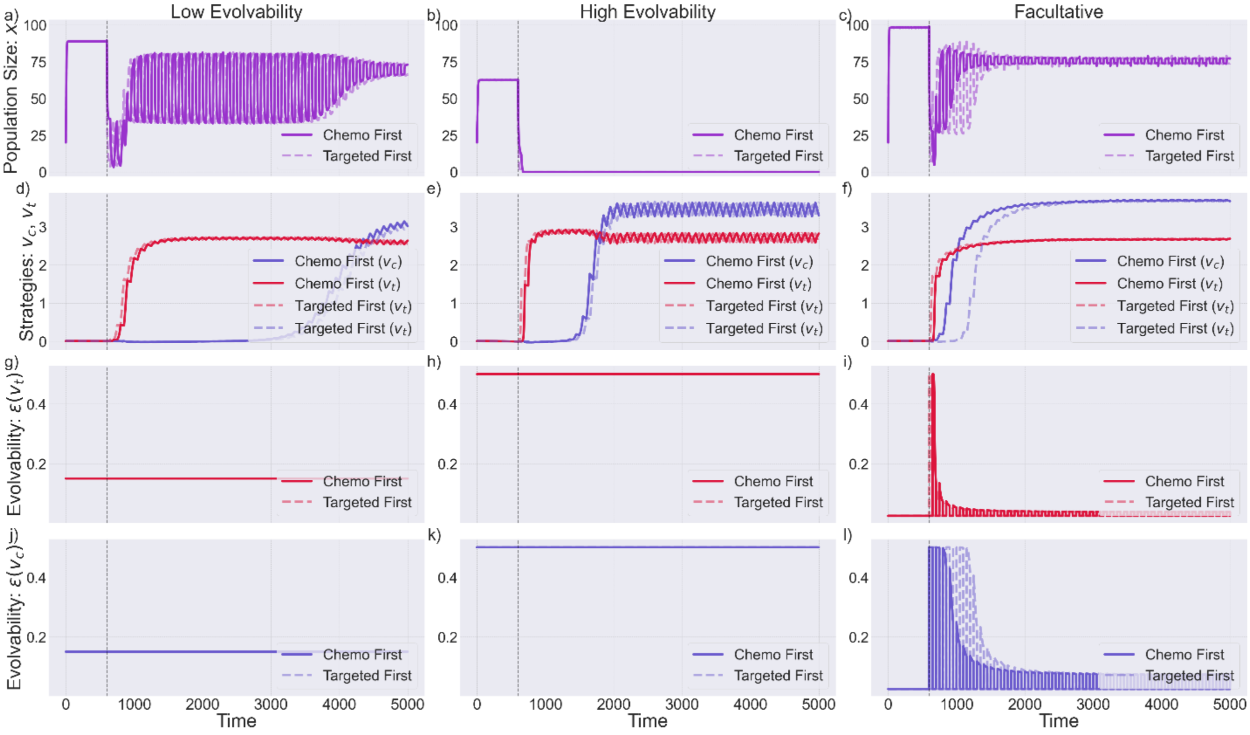


Figure S3: Comparison of dynamics under double bind therapy for alternative case: Population dynamics (a,b,c), resistance strategy dynamics (d,e,f), and evolvability (g-l) dynamics of the slow-evolving, fast-evolving, and facultative cancer population under double bind therapy. Therapy is initiated at $t = 600$. Solid lines represent the regimen where chemotherapy is given first, followed by targeted therapy; dashed lines represent the reverse order. The cycle duration is **short**, i.e., therapy is switched every **50** steps. Parameters used are shown in Table S1.

Alternative Case: Double Bind

Cycle Duration: 200

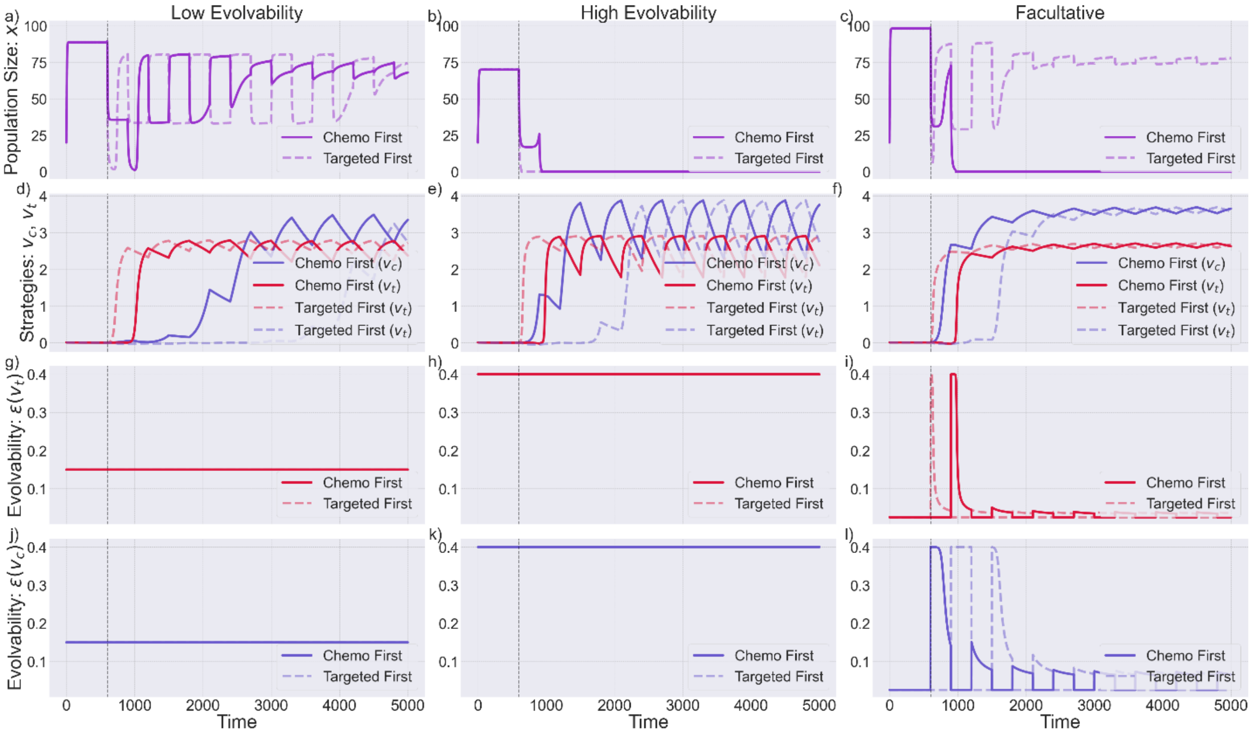


Figure S4: Comparison of dynamics under double bind therapy for alternative case: Population dynamics (a,b,c), resistance strategy dynamics (d,e,f), and evolvability (g-l) dynamics of the slow-evolving, fast-evolving, and facultative cancer population under double bind therapy. Therapy is initiated at $t = 600$. Solid lines represent the regimen where chemotherapy is given first, followed by targeted therapy; dashed lines represent the reverse order. The cycle duration is **long**, i.e., therapy is switched every **300** steps. Parameters used are shown in Table S1.

Alternative Case: Combination Therapy

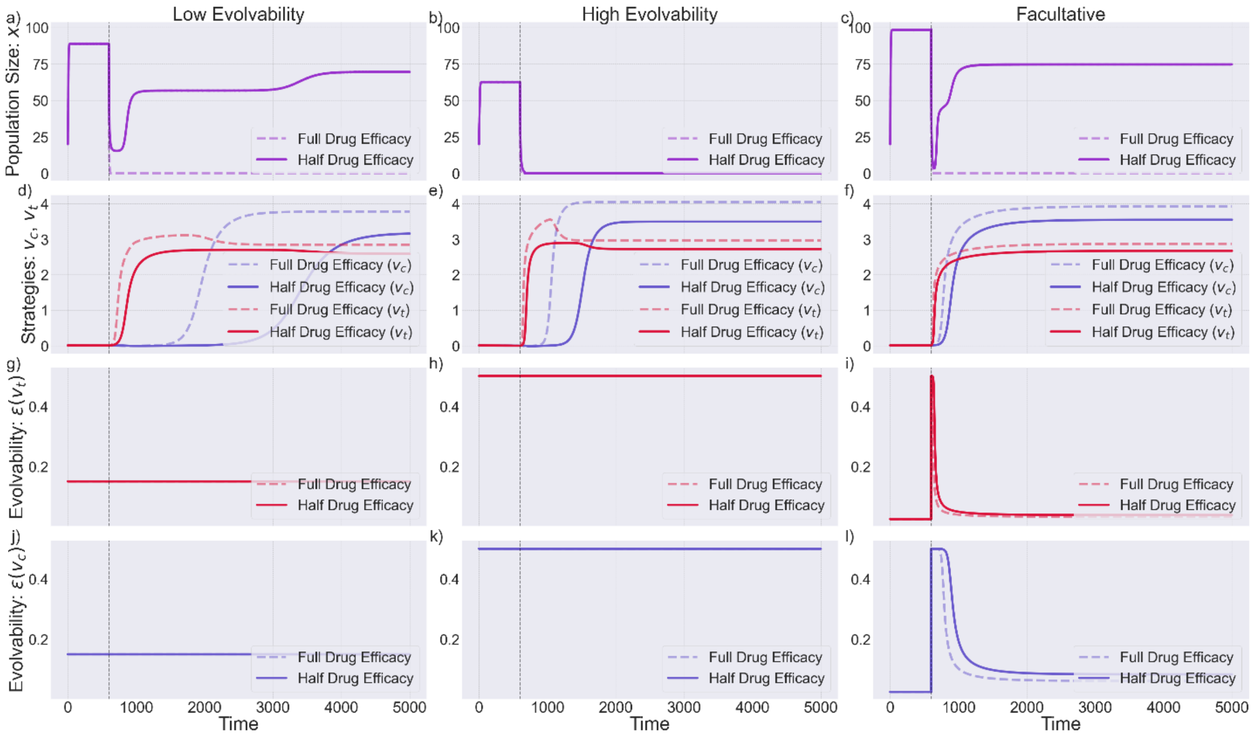


Figure S5: Comparison of dynamics under combination therapy for alternative case: Population dynamics (a,b,c), resistance strategy dynamics (d,e,f), and evolvability (g-l) dynamics of the slow-evolving, fast-evolving, and facultative cancer population under combination therapy. Therapy is initiated at $t = 600$. Solid lines represent the regimen where the full drug dose is administered; dashed lines show the regimen where half of the drug dose is administered to reduce toxicity. Parameters used are shown in Table 1 & 2.

Alternative Case: Double Bind

Cycle Duration: 500

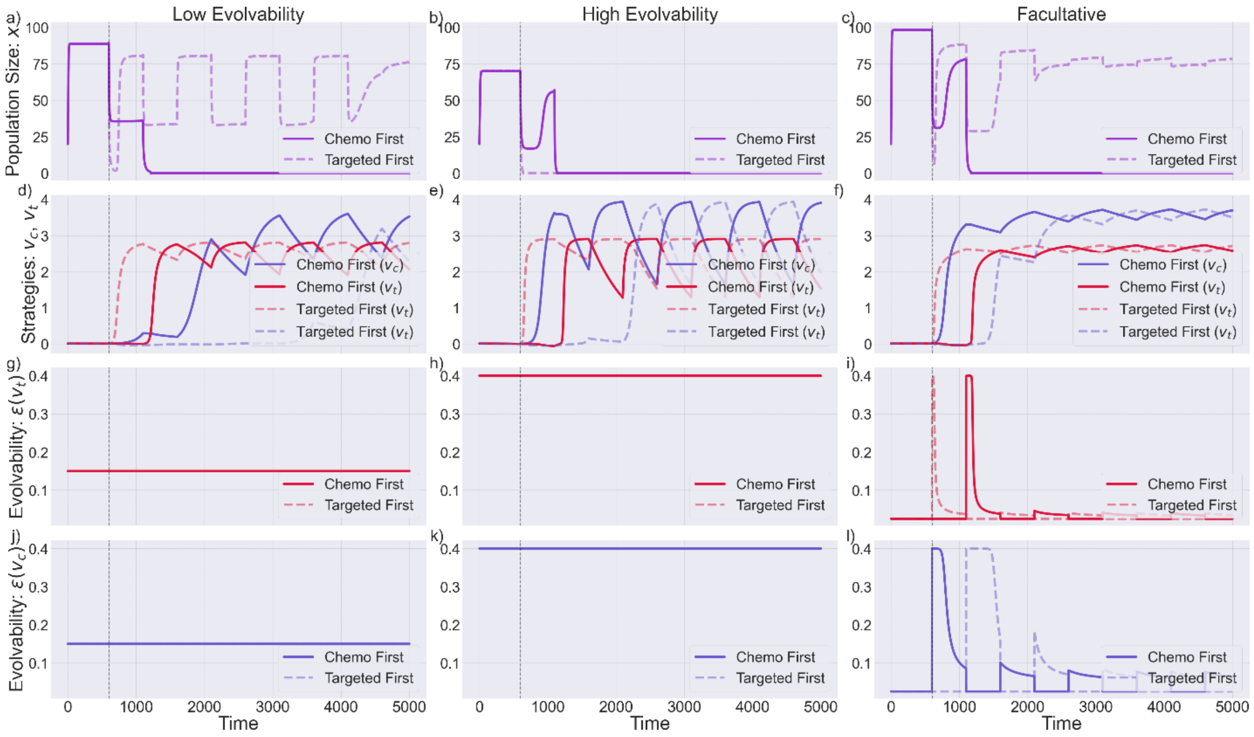


Figure S6: Comparison of dynamics under double bind therapy for alternative case: Population dynamics (a,b,c), resistance strategy dynamics (d,e,f), and evolvability (g-l) dynamics of the slow-evolving, fast-evolving, and facultative cancer population under double bind therapy. Therapy is initiated at $t = 600$. Solid lines represent the regimen where chemotherapy is given first, followed by targeted therapy; dashed lines represent the reverse order. In this case therapy is switched every **500** steps. Parameters used are shown in Table S1.
